# Modeling cardiac microcirculation for the simulation of coronary flow and 3D myocardial perfusion

**DOI:** 10.1101/2024.04.11.588994

**Authors:** Giovanni Montino Pelagi, Francesco Regazzoni, Jacques M. Huyghe, Andrea Baggiano, Marco Alì, Silvia Bertoluzza, Giovanni Valbusa, Gianluca Pontone, Christian Vergara

**Affiliations:** LABS, Dipartimento di Chimica, Materiali e Ingegneria Chimica “Giulio Natta”, Politecnico di Milano, Piazza Leonardo da Vinci 32, Milan, 20133, Italy; MOX, Dipartimento di Matematica, Politecnico di Milano, Piazza Leonardo da Vinci 32, Milan, 20133, Italy; School of Engineering, University of Limerick, Limerick V94 T9PX, Limerick, Ireland; Eindhoven University of Technology, 5600 MB, Eindhoven, Netherlands; Perioperative Cardiology and Cardiovascular Imaging Department, Centro Cardiologico Monzino IRCCS, Via Carlo Parea 4, Milan, 20138, Italy; Department of Clinical Sciences and Community Health, University of Milan, Milan, Italy; Bracco Imaging S.p.A., Via Caduti di Marcinelle 13, Milan, 20134, Italy; Department of Diagnostic Imaging and Stereotactic Radiosurgery, Centro Diagnostico Italiano S.p.A. Via Saint Bon 20, Milan, 20147, Italy; IMATI, CNR, Pavia, Italy; Department of Biomedical, Surgical and Dental Sciences, University of Milan, Milan, 20134, Italy

**Keywords:** coronary artery disease, fractional flow reserve, myocardial perfusion, myocardial blood flow, computational modeling, coronary pressure

## Abstract

**Purpose:** accurate modeling of blood dynamics in the coronary microcirculation is a crucial step towards the clinical application of *in silico* methods for the diagnosis of coronary artery disease (CAD). In this work, we present a new mathematical model of microcirculatory hemodynamics accounting for microvasculature compliance and cardiac contraction; we also present its application to a full simulation of hyperemic coronary blood flow and 3D myocardial perfusion in real clinical cases.

**Methods:** microvasculature hemodynamics is modeled with a *compliant* multi-compartment Darcy formulation, with the new compliance terms depending on the local intramyocardial pressure generated by cardiac contraction. Nonlinear analytical relationships for vessels distensibility are included based on experimental data, and all the parameters of the model are reformulated based on histologically relevant quantities, allowing a deeper model personalization.

**Results:** Phasic flow patterns of high arterial inflow in diastole and venous outflow in systole are obtained, with flow waveforms morphology and pressure distribution along the microcirculation reproduced in accordance with experimental and *in vivo* measures. Phasic diameter change for arterioles and capillaries is also obtained with relevant differences depending on the depth location. Coronary blood dynamics exhibits a disturbed flow at the systolic onset, while the obtained 3D perfusion maps reproduce the systolic impediment effect and show relevant regional and transmural heterogeneities in myocardial blood flow (MBF).

**Conclusion:** the proposed model successfully reproduces microvasculature hemodynamics over the whole heartbeat and along the entire intramural vessels. Quantification of phasic flow patterns, diameter changes, regional and transmural heterogeneities in MBF represent key steps ahead in the direction of the predictive simulation of cardiac perfusion.

## 1 Introduction

Cardiac blood perfusion is the central physiological process that guarantees the metabolic sustenance of the heart muscle, requiring a dedicated circulatory system known as the coronary circulation. Defective perfusion is generally caused by a narrowing or blockage of a coronary artery, a condition known as Coronary Artery Disease (CAD), and leads to major consequences such as myocardial ischemia, infarction and heart failure. Large clinical studies have shown that the combined knowledge of pressure drop in the large coronaries and Myocardial Blood Flow (MBF) at the tissue level leads to the best management of patients suffering from CAD [1, 2]. At present, however, this knowledge can be achieved only through multiple imaging exams, often including radiation exposure and potentially invasive procedures [3].

In this context, mathematical models and computational simulations of coronary hemodynamics integrating blood flow in the large coronary arteries and myocardial perfusion hold great potential to provide clinically relevant information, especially when tailored to a specific subject using *in vivo* radiological images. Still, an accurate mathematical description of coronary hemodynamics remains a challenge because of two main reasons: firstly, the coronary circulation spans over a broad range of length scales (from few millimeters to few microns of vessel diameter), making it impossible to run even 1D fluid-dynamics simulations in the fully resolved tree; secondly, cardiac contraction deeply affects coronary flow, mainly through the well known systolic impediment effect [4], which is challenging to model in an effective way. To address the first issue, previous works have proposed either a focus on large coronaries with outflow conditions, surrogating microvasculature, based on lumped parameter models [5, 6] or on extendend Murray’s law [7]; or multiscale models, often treating blood dynamics in the microcirculation through a homogenized porous medium approach (Darcy equations, [8]), coupled with a 1D [9] or 3D [10] description of fluid-dynamics in the large coronaries. This has been further extended with the proposal of multicompartment Darcy formulations to account for the different length scales in the microcirculation [11–14].

To cope with cardiac contraction, previous approaches relied on poromechanics as a way to model flow through a saturated porous medium subjected to mechanical activation [15–17], possibly coupled with coronary arterial networks [18–20]. In a previous work [21], we proposed an *effective* inlet pressure condition for the large coronaries to surrogate the effects of cardiac contraction in a multiscale coupled model of cardiac perfusion. However, the first models are difficult to personalize and have never been applied to real clinical scenarios, whereas what proposed in [21] does not provide a sufficient accuracy for the distribution of blood flow at the tissue level.

In this work, to overcome these limitations, we start from the multiscale perfusion model presented in [13] and we propose:

1. A new mathematical formulation of the multicompartment Darcy model to account for cardiac mechanics and microvasculature compliance;
2. A new, data-driven calibration of the Darcy model parameters in view of simulations of hyperemic coronary flow in real clinical cases.

This new model for microvascular hemodynamics is coupled with 3D fluid-dynamics equations for the blood dynamics in the large coronaries to run an integrated analysis (at all levels of the coronary tree) in subjects whose heart and coronary geometries have been reconstructed from *in vivo* CT images.

To the best of our knowledge, this is the first computational model, incorporating details along all the coronary tree and effects of cardiac contraction on perfusion, which has been calibrated for an application to real clinical cases. We believe that this work is a crucial step towards a predictive application of perfusion modeling in a clinical setting and the use of computational methods to reproduce functional imaging of stress Computed Tomographic Perfusion (stress-CTP) [1].

## 2. Methods

In Section 2.1 we introduce the new mathematical formulation for the Darcy multi-compartment model in a domain subjected to a cyclic mechanical stress caused by cardiac contraction. In Sections 2.2 and 2.3 we provide an overview of the new parameters introduced and we propose a data-driven approach for their calibration, whereas in Section 2.4 the specific choice to surrogate the effects of cardiac contraction is presented. Section 2.5 includes the details regarding the discretization and numerical solution of the resulting mathematical problem, while in Section 2.7 the general setup used for the numerical simulations is described.

### 2.1 Model of microcirculation hemodynamics

When the coronary arteries penetrate the myocardial surface, they progressively branch into smaller vessels in a tree-like structure known as the *intramural* circulation. Given the huge number of vessels, a homogenized approach where hemodynamics is described as a flow through a porous medium is well suited to describe hemodynamics in this part of the coronary tree [8, 15, 22].

To account for the different length scales (diameters from *d* ≃ 5 *μ*m, capillaries, up to *d* ≃ 500 *μ*m, small arteries) as well as for the mechanical activity of the heart, we start from the three-compartment primal Darcy formulation presented in [8, 13] and we generalize it with the addition of a compliance term resulting from vessels distensibility. For the compartments, we consider the following subdivision: small arteries (comp. 1, *d* between 100 and 500 *μ*m), arterioles (comp. 2, *d* between 8 and 100 *μ*m), capillaries (comp. 3, *d* between 4 and 8 *μ*m). The strong formulation for a generic compartment *i* = 1, 2, 3 reads:

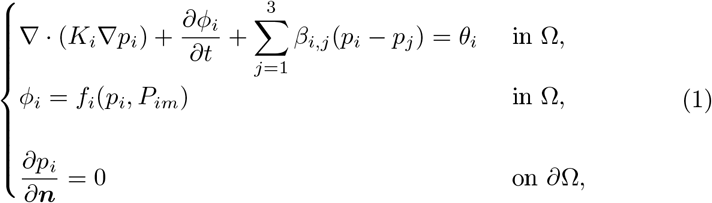

where, for each compartment *i, p*_*i*_ is the unknown intraluminal blood pressure, *K*_*i*_ is the permeability, *β*_*i,j*_ is the mass exchange coefficient with compartment *j*; *θ*_*i*_ is a distributed mass source/sink term accounting both for the mass source in compartment 1 that represents the flow coming from the large arteries (for example provided by the solution of a Navier-Stokes problem, see [13, 14] and Section 2.7) and for the mass sink in compartment 3 representing the venous return, i.e. *θ*_3_ = −*γ*(*p*_3_ − *p*_*ra*_), *p*_*ra*_ being the right atrium pressure. Notice that the equations related to each compartment are solved in the same computational domain Ω, that is the left ventricular free wall reported in Figure 1, top-left, meaning that we assume each compartment of intramural vessels to coexist in the same volume.

**Figure 1:**
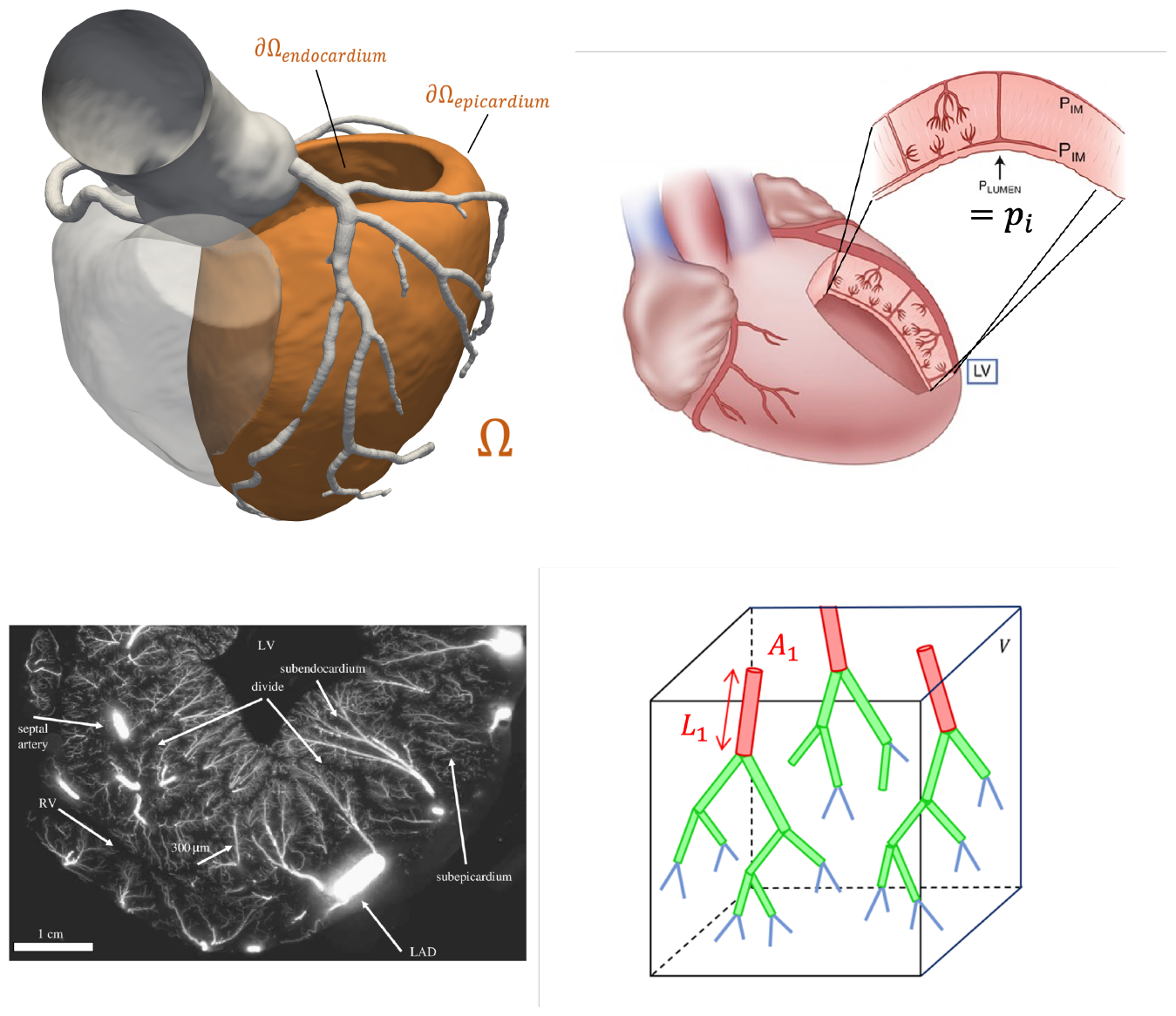
Top-left: computational domain Ω (consisting of the left ventricular free wall, in orange) for the microcirculation model. Aortic root, epicardial coronary tree and right ventricular chamber (shaded) are also displayed but not a part of Ω. Top-right: schematic representation of the co-existence of intramural vessels and myocardial tissue within the left ventricular free wall (adapted from [23]). Bottom-left: *ex vivo* cryomicrotome image of coronary intramural circulation showing the organization of vessels within the ventricular wall (reproduced from [24]). Bottom-right: schematic visualization of the hierarchical organization of the intramural network within a homogenization volume *V* : coloring represents the belonging of a vessel to a specific class (i.e. small arteries, arterioles or capillaries) which corresponds to a specific Darcy compartment in our model. Vessels cross section *A*_*i*_ and average length *L*_*i*_ are indicated for the first compartment, with vessels density 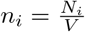 for each compartment *i*.

The new compliance term 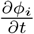 represents the time variation of the fluid volume fraction *φ*_*i*_ (i.e. the porosity of the *i th* compartment), which we model with a suitable set of functions *f*_*i*_ representing the relationship between compartment porosity, the intraluminal pressure *p*_*i*_ and the given intramyocardial pressure *P*_*im*_, generated within the cardiac tissue by the heart contraction. A schematic representation of the intraluminal/extraluminal spaces with their pressures is reported in Figure 1, top-right, while in Sections 2.4 we propose a specific treatment for the computation of *P*_*im*_, with the main modeling assumption being that *P*_*im*_ is considered independent of the intraluminal blood pressure and prescribed as a given datum.

Considering the Darcy compartments as networks of cylindrical vessels (as schematized in Figure 1, bottom-right), the local fluid volume fraction of compartment *i* in a given homogenization volume *V* can be written, by definition, as:

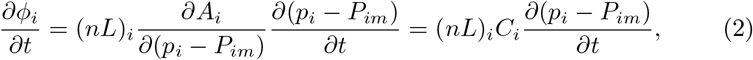

where *N*_*i*_ is the total number of vessels belonging to compartment *i, L*_*i*_ is the average vessel length and *A*_*i*_ is the average vessel cross section. The ratio 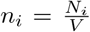 is the local density of vessels belonging to compartment *i*, whereas the product (*nL*)_*i*_ is often denoted as the vessel *length density*. Given that the intramural vessels are mainly aligned in a transmural direction (i.e. in a direction perpendicular to the myocardial fibrils, see Figure 1, bottom-left and [24]) we assume that cardiac contraction affects exclusively the cross sectional area, whereas the vessels density and length remain constant. Further, we assume that vessels cross section explicitly depend on the transluminal pressure difference (*p*_*i*_ −*P*_*im*_), as supported by experimental measures [25]. Under these hypotheses, the time derivative of the fluid volume fraction becomes:

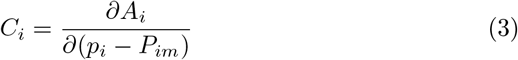

Where

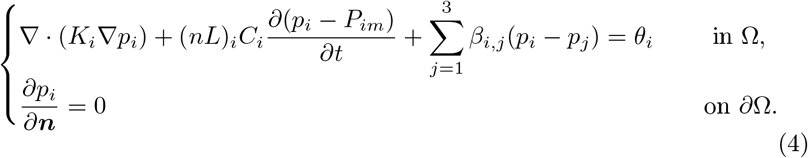

represents the *distensibility* of vessels in compartment *i*, which is, in general, dependent on the transluminal pressure difference (*p*_*i*_ −*P*_*im*_).

By substituting eq. (2) into (1), we obtain the final formulation of the *compliant* multicompartment Darcy model:

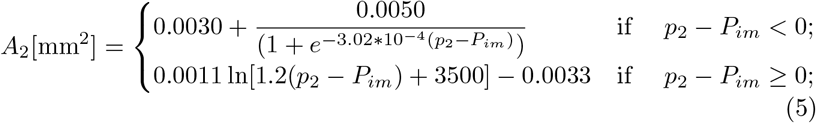

for *i* = 1, 2, 3. Notice that parameters *K*_*i*_, *β*_*i,j*_ and *C*_*i*_ in (4) are dependent on the the transluminal pressure difference (*p*_*i*_ −*P*_*im*_) and are computed using the methods exposed in the following two sections.

### 2.2 Constitutive relations for vessel distensibility

As seen by its definition (3), vessels distensibility *C*_*i*_ represents the variation of cross-sectional area with respect to variations in transluminal pressure difference. Since the histological structure of coronary microvessels changes depending on their diameter, the constitutive relationships we used for *C*_*i*_ are compartment-specific and their choice is driven by experimental data. It is important to note that, since we are interested in hyperemic coronary flow, we used data related only to a vessel condition of maximal vasodilation, ruling out the effects of vascular tone and autoregulation mechanisms. This is motivated by the fact that, at maximal hyperemia, these mechanisms are exhausted and the vessel wall can be modeled as a fully passive structure.

For the compartment-specific constitutive relationships we adopt the consider what follows:

1. Small arteries (comp. 1) have a relatively thick wall structure consisting of collagen and smooth muscle cells, thus we consider these vessels as rigid and we set

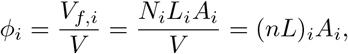

where the specific value chosen for *A*_1_ corresponds to a diameter *d* = 150*μm*, which we consider as mean diameter for vessels of this class.
2. Arterioles (comp. 2) have been found to be much more distensible than the small arteries during *in vivo* observations in animal experiments [26, 27]. Also, experimental measures of arterioles have shown a highly nonlinear relationship between transluminal pressure and cross sectional area [25]. To capture this behaviour, we fitted a logarithmic relationship on data from isolated arterioles [25], see Figure 2a. Since we have at disposal data only for *p*_2_ −*P*_*im*_ *>* 0, we extend this curve through a sigmoid function in the negative region. The analytical expression is given by:

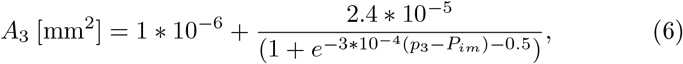

so that from (3) we obtain

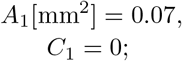
3. Capillaries (comp. 3) do not have any muscle cells, having only a single layer of endothelial cells. However, experimental observations have shown that they are surprisingly resistant to systolic compression, exhibiting a relatively low change in diameter between the diastolic (*d*≃ 5.4 *μ*m) and systolic (*d*≃ 4.3 *μ*m) phases [28]. Differently from the case of arterioles, there are no experimental data relating capillary cross section and transluminal pressure. We assume as a modeling abstraction a curve for the capillary cross section built from the values of systolic/diastolic diameter in [28] with a sigmoid shape similar to *A*_2_, see Figure 2b. Its analytical expression is given by:

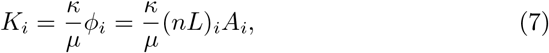

so that we obtain

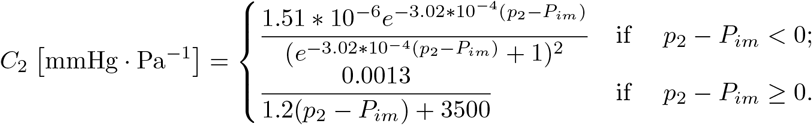

**Figure 2:**
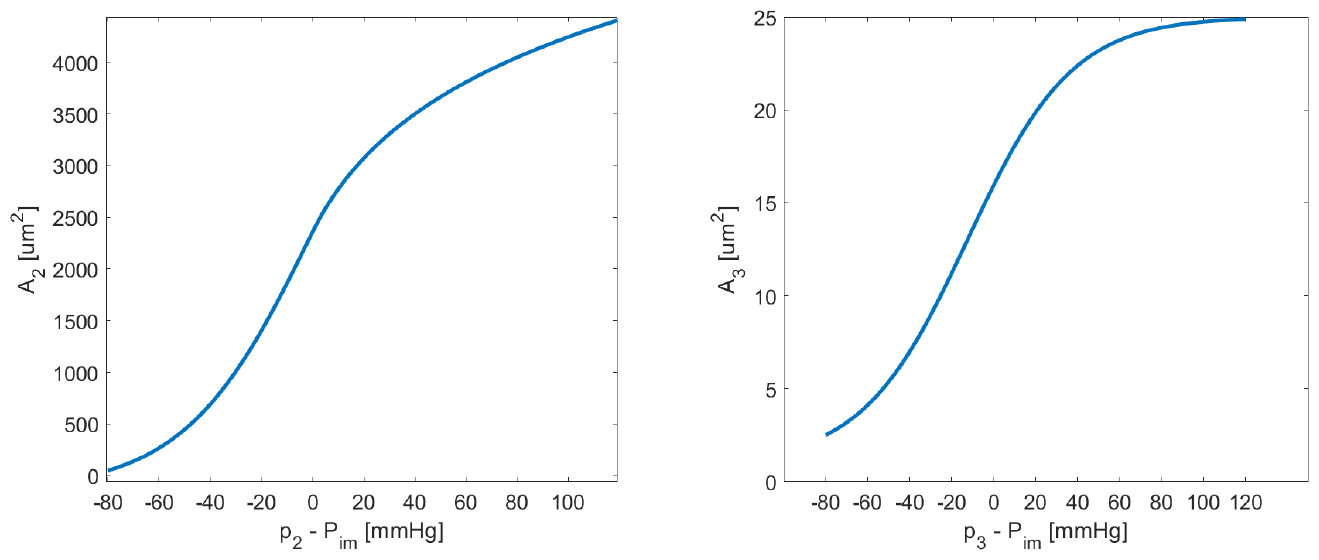
Constitutive curves relating vessels cross section and transluminal pressure for arterioles (left) and capillaries (right).

### 2.3 Estimation of Darcy parameters

For permeabilities *K*_*i*_ we assume a direct proportionality with the porosity, which coincides with the fluid volume fraction (the porous medium is saturated):

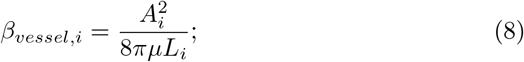

where *κ* is the specific permeability of the fluid/matrix system, considered as independent from the Darcy compartment, and *μ* is the blood dynamic viscosity; see the Discussion section for such choices.

The conductance coefficients *β*_*i,j*_ mediate mass transfer between compartments, so for their formulation we start considering each vessel as a Poiseuille-like resistor with conductance [29]:

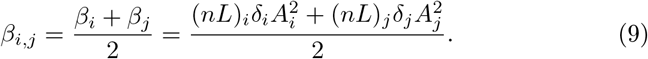

then we extend this formula to the whole compartment, by considering it as a network of conductances in series and parallel, which is motivated by the tree-like structure of vessels networks. The whole network conductance is then proportional to the one of the single vessels and to the length density:

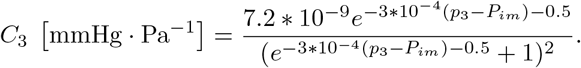

Thus, exploiting (8), we obtain:

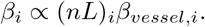

where *δ*_*i*_ is a coefficient taking into account the morphometry of the network and includes the blood viscosity. Since we can assume that mass exchanges between compartments depend on the conductances of both the upstream and downstream networks, we finally take as inter-compartment conductance *β*_*i,j*_ the expression:

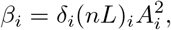

Parameters *κ* in eq. (7) and *δ*_*i*_ in eq. (9) are difficult to estimate from data, so for their computation we rely on a calibration procedure exposed in Section 2.7, alongside a list of all the other parameters used in the simulations (see Table 1).

**Table 1:**
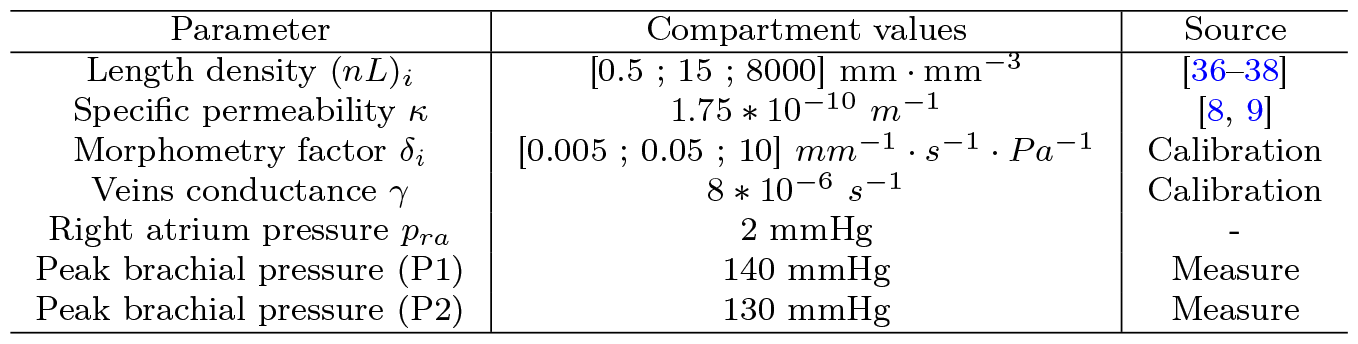
List of parameters used for the simulations. Calibration refers to a trial-and-errors procedure on patient P1 to recover experimental data.

### 2.4 Estimation of the intramyocardial pressure

The knowledge of the intramyocardial pressure *P*_*im*_ is fundamental to build curves *A*_*i*_ and *C*_*i*_ as well as the compliance term 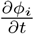 in (1). It allows to model the effect of cardiac mechanics on microcirculation hemodynamics. According to previous studies [30], most of the experimental observations related to coronary hemodynamics can be explained by considering cyclic changes in *P*_*im*_ induced by and closely following the pressure *P*_*LV*_ generated inside the ventricular chamber. Other mechanisms, such as the shortening-induced intracellular pressure, were found to play a role only in the case of specific states of altered contractility.

Given these findings, we consider for *P*_*im*_ the temporal waveform of *P*_*LV*_ obtained from an electromechanics simulation [31]), that we report in Figure 3-left, which includes also a comparison with the aortic pressure. Since experimental findings pointed out that intramyocardial pressure decreases almost linearly from the subendocardium to the subepicardium [32], we include a linear transmural modulation of *P*_*im*_ such that:

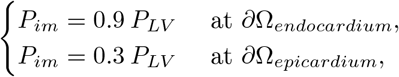

which is in accordance with the quantitative experimental data in [32, 33]. A representation of this transmural modulation is reported at the systolic peak in Figure 3-middle, together with a left ventricular distribution, see Figure 3-right .

**Figure 3:**
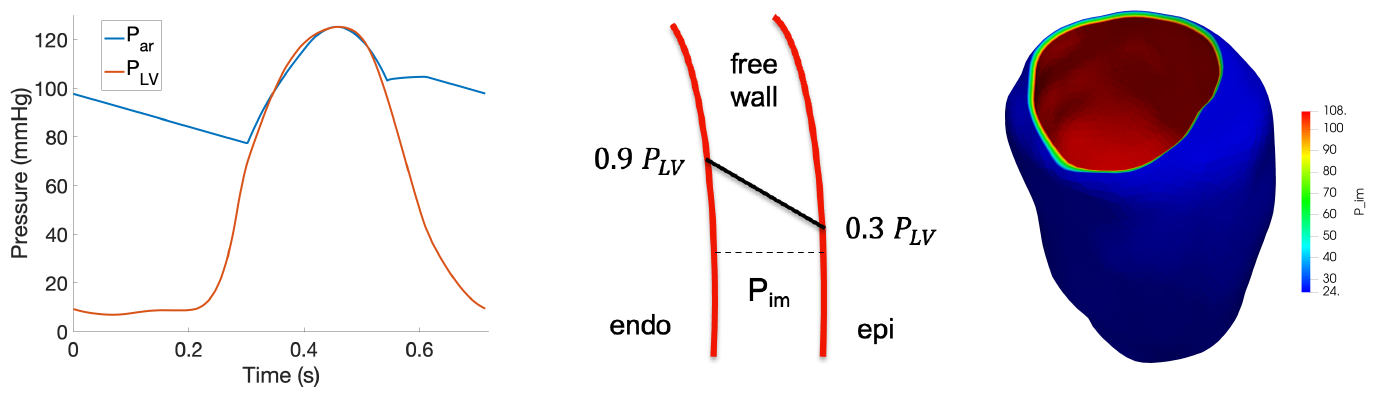
Left: time waveform of the pressure in the left ventricular chamber, obtained with an electromechanics simulation [31]. Middle: transmural modulation of intramyocardial pressure *P*_*im*_ in the left ventricular free wall. Right: 3D representation of the transmural modulation displayed at systolic peak.

### 2.5 Numerical discretization

Parameters *K*_*i*_, *β*_*i,j*_ and the compliance term *C*_*i*_, computed with the datadriven approach described in Sections 2.2-2.3, introduce significant nonlinearities in the compliant multicompartment Darcy model (4). To cope with this issue, we rely on a linearized version of (4) obtained with a first-order finite difference discretization for the time derivatives and a semi-implicit treatment for the unknowns. Given a function *v*(*t*), we introduce a partition of time domain based on discrete instants *t*^*n*^ = *n*Δ*t, n* = 0, 1, …, with Δ*t* being the time discretization parameter, and we denote the approximated quantity as *v*^*n*^≃ *v*(*t*^*n*^).

Accordingly, the time-discretized model reads:

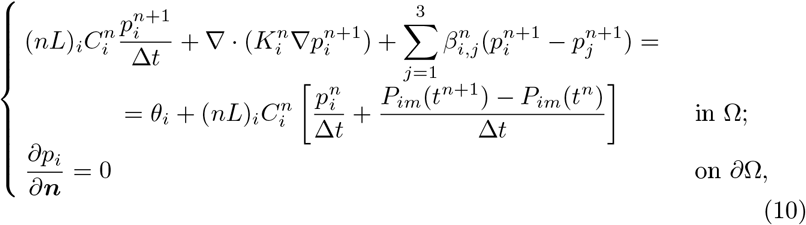

where *P*_*im*_ is the given intramyocardial pressure.

### 2.6 Coupling with epicardial coronaries

The above Darcy problem is coupled with large coronaries hemodynamics, where a 3D fluid-dynamics problem given by the incompressible Newtonian Navier-Stokes equations is considered [13]. For the time discretization, a first order semi-implicit time scheme is employed. The fluid mesh, characterized by a value of *h* which is proportional to the lumen diameter with an average value of 0.4 mm, is obtained after convergence tests for the computational domain reported in Figure 4-left. At the two coronary inlets 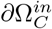 we prescribe patient-specific pressure waveforms.

**Figure 4:**
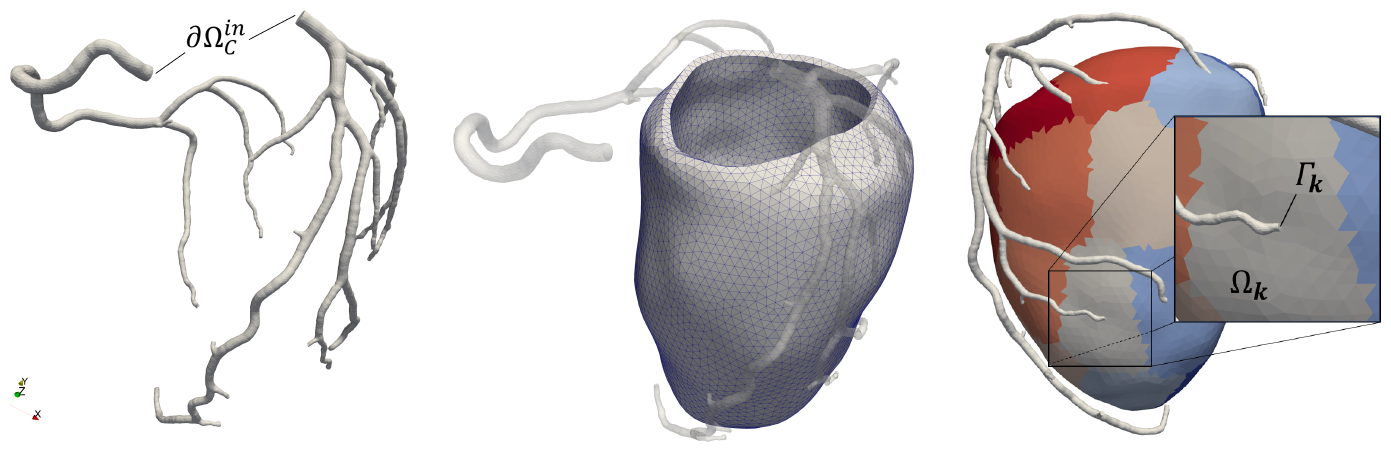
Left: segmented domain used for the solution of the 3D hemodynamics problem. Middle: segmented domain, with mesh detail, used for the solution of the multi-compartment Darcy problem. Right: representation of the coupling coupling between the two problems: each coronary outlet Γ_*k*_ is coupled with a corresponding perfusion territory Ω_*k*_.

Downstream, the new multicompartment Darcy fomulation reported in Section 2.5 and representing microcirculation hemodynamics is solved in the left ventricular free wall (Figure 4-middle). To couple the two subproblems, each coronary outlet is associated with a perfusion territory in the myocardium, see Figure 4-right. Interface conditions representing force balance are prescribed at each coronary outlet based on the mean pressure of Darcy compartment 1 in the corresponding perfusion territory (see [13] for the mathematical details). Interface conditions representing mass conservation are prescribed in the Darcy problem through the following source terms *θ*_*i*_:

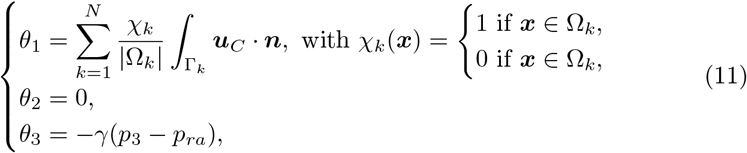

where *N* is the total number of coronary outlets. In the first of (11), which represents the mass source in the first Darcy compartment, ***u***_*C*_ is the coronary velocity unknown and the term 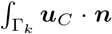 is the flow through the coronary outlet Γ_*k*_, |Ω_*k*_ | is the volume of the corresponding perfusion territory, and *χ*_*k*_ is a characteristic function ensuring that each perfusion territory receives blood only from the corresponding coronary outlet Γ_*k*_. Notice also the third line of (11), representing the mass sink in the third Darcy compartment due to the venous return and featuring the constant parameters *γ* (conductance of the whole coronary venous circulation) and *p*_*ra*_ (right atrium pressure), whose values are reported in Table 1.

### 2.7 Simulation setup

Coronary Blood Flow (CBF) and perfusion simulations in hyperemic conditions (CBF-Perfusion simulations) are carried out for two patients from Centro Cardiologico Monzino. These patients were chosen among subjects with the following characteristics:

1. No history of previous major cardiac adverse events;
2. Symptoms of coronary artery disease including *angina pectoris*;
3. No actual anatomical signs of coronary artery disease (ruled out by imaging exams), including coronary stenoses and atherosclerotic plaques.

These patients are therefore representative of a population of high-risk, symptomatic subjects that, however, have shown no sign of CAD or myocardial ischemia.

For both patients, the left ventricular and coronary geometries were segmented from contrast-enhanced coronary Computed Tomographic Angiography (cCTA) images and meshed using the software packages VMTK [34]. An example of the computational domains obtained is reported in Figure 4.

Aortic root pressure curves for the inlet boundary conditions for the Navier-Stokes equations in the epicardial coronaries were built in a personalized way from basic measures [21], whereas the left ventricular pressure *P*_*LV*_ curves, used to build *P*_*im*_ were tailored, starting from the base waveform of Figure 3 (obtained from an electromechanics simulation [31]), to match the patient’s systolic interval and peak aortic pressure, see Table 1 and methods in [21]. Time discretization parameter Δ*t* is set equal to 2 *ms*.

Table 1 reports a list of the values for all the parameters used alongside an indication on how they were chosen. In particular, parameters *κ, γ* and *δ*_*i*_ were calibrated o a single patient (patient P1) with two targets:

1. Reproduce a distribution of pressure, along the microvasculature, matching experimental data from [35];
2. Reproduce an in-space average MBF, taken as the time-averaged capillary flow, matching the value obtained from the stress-CTP exam.

The same parameters were then used also for another patient P2, that represents, therefore, a validation case.

All the simulations were run using the software life^x^, a high performance library for Finite Elements simulations of multiphysics, multiscale and multidomain problems developed at MOX - Dipartimento di Matematica, in cooperation with LaBS - Dipartimento di Chimica, Materiali e Ingegneria Chimica, both at Politecnico di Milano [39, 40].

## 3 Results

Simulations results are analyzed in Section 3.1 in terms of the following outputs:

1. Time evolution of the epicardial coronary flow, computed from the Navier-Stokes (NS) model, and of the venous outflow computed from the Darcy model;
2. Time evolution of the in-space average pressure within the Darcy compartments (intramural blood pressure);
3. Time evolution of arteriolar flow, capillary flow and vessel diameter at the subendocardium, midmyocardium and subepicardium, computed from the Darcy model;
4. 3D distribution of capillary flow (from Darcy) and 3D velocity patterns in the epicardial arteries (from NS) along the whole heartbeat.

All the analyzed outputs are compared with *in vivo* human measurement or experimental data, when available. In Section 3.2 we report a sensitivity analysis with respect to the new parameters introduced with the proposed model.

### 3.1 Analysis of hemodynamics results

Figure 5, left-top and left-middle, reports the time evolution of the arterial inflows and venous outflow, the latter computed as the in-space average of *θ*_3_ as defined in (11), for patients P1 and P2. The physiological phasic pattern of high diastolic inflow and high systolic outflow can be clearly seen; moreover, we find an excellent accordance of arterial waveforms with *in vivo* experimental measures reported in [41, 42] and depicted in Figure 5, left-bottom.

**Figure 5:**
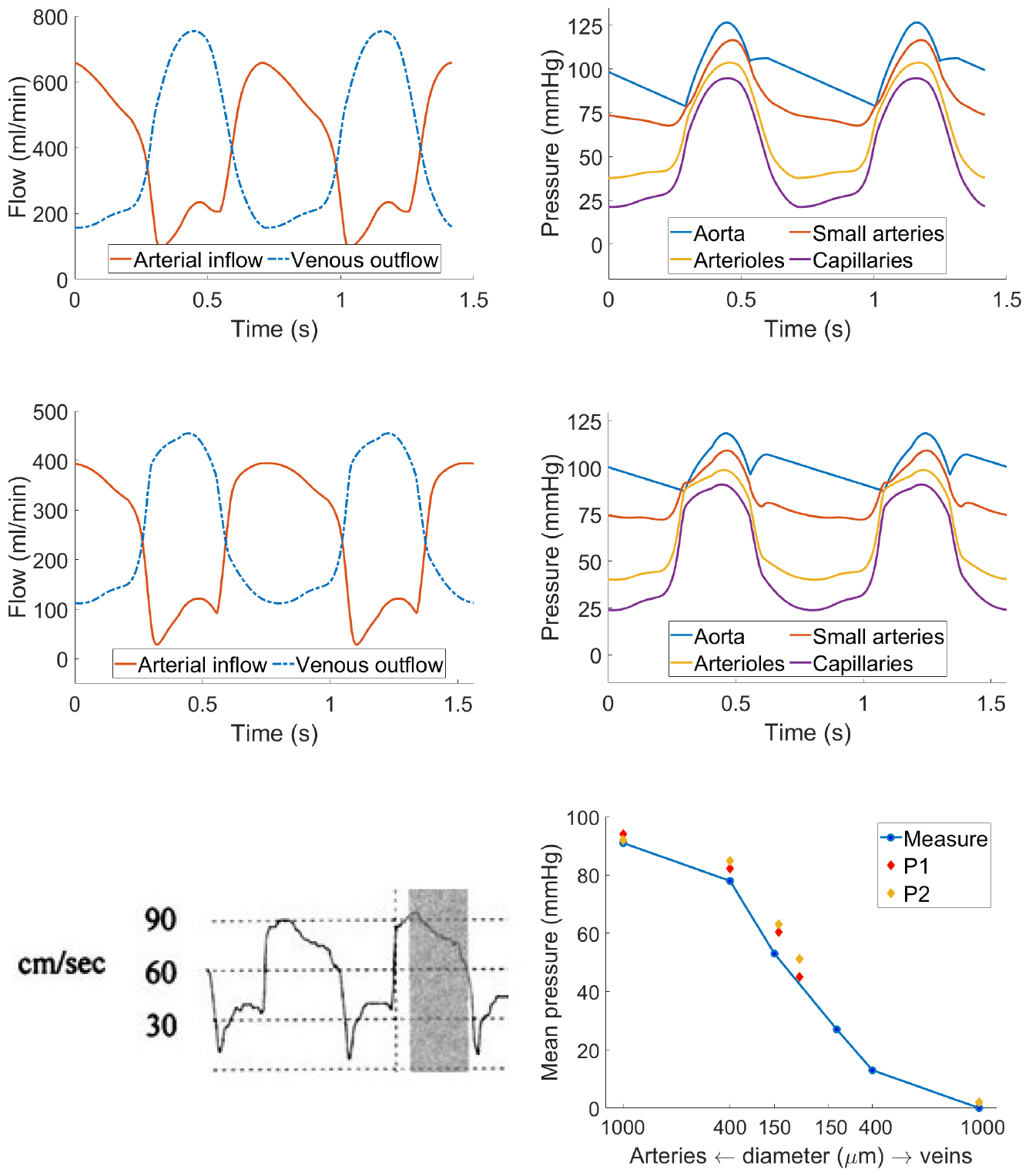
Left: total left arterial inflow and venous outflow over time for patient P1 (top) and P2 (middle) as compared with a Doppler intracoronary velocity measure reproduced from [41] (bottom). Right: average-in-space pressure in the Darcy compartment and aortic pressure over time for patient P1 (top) and P2 (middle). Right-bottom: averaged (both in space and in time) pressure values for both patients compared to experimental measures reported in [35]. Straight blue lines only for visualization purposes.

Table 2 summarizes some relevant quantities regarding the morphology of the flow waveform as compared with the *in vivo* Doppler measurements in rest conditions reported in [43]. We report an excellent agreement in terms of the systolic/diastolic flow ratio in the Left Anterior Descending (LAD) artery, whereas this ratio is underestimated in the case of the Right Coronary Artery (RCA). This is in agreement with our geometric model choices, which do not include the RCA branches perfusing the right ventricle, where the flow follows the aortic pressure waveform, thus featuring its peak during systole. Our computed peak/mean flow ratios show a slight underestimation compared to the measures, which could be due to small mismatches in the timing between the aortic and intramyocardial pressure waveforms used as input, resulting in a smoothed early diastolic flow peak.

**Table 2:** List of flow-related ratios computed by our model as compared to the *in vivo* Doppler measures reported in [43].

| Ratio | P1: LAD-RCA | P2: LAD-RCA | Measure: LAD-RCA |
| --- | --- | --- | --- |
| Systolic / Diastolic peak flow ratio | 0.35 - 0.38 | 0.31 - 0.31 | 0.37 - 0.97 |
| Systolic / Diastolic mean flow ratio | 0.35 - 0.39 | 0.28 - 0.29 | 0.22 - 0.85 |
| Mean flow / Peak flow (Systole) | 0.45 - 0.48 | 0.75 - 0.74 | 0.32 - 0.38 |
| Mean flow / Peak flow (Diastole) | 0.75 - 0.78 | 0.81 - 0.81 | 0.57 - 0.46 |

Figure 5, right-top and right-middle, reports the time evolution of the (inspace averaged) Darcy pressure for all the compartments, compared with the aortic pressure curve for patients P1 and P2. We can see that only small arteries follow the waveform of the aortic pressure, whereas arteriolar pressure is dominated by the intramyocardial pressure generated by contraction, with increasing pressure during diastole likely caused by the vessel filling with blood.

The curves show blood pressurization due to contraction in systole. We report an in-time mean value of the Darcy pressures of 82.2, 60.4, and 45.0 *mmHg* (patient P1) and 84.9, 63.1 and 51.3 *mmHg* (patient P2) for the small arteries, arterioles and capillaries respectively. All these findings reproduce what experimentally measured and reported in [35] and depicted in Figure 5, bottom-right. Notably, we did not observe any retrograde flow in the early systole, whose absence may be due to the hyperemic conditions.

Figure 6-top reports the arteriolar and capillary blood flow waveforms over time (patient P1), both computed at three sample points in the mid-anterior left ventricular wall located at different depths: subepicardium (1 *mm* below epicardial surface), midmyocardium (in the middle), and subendocardium (1 *mm* below endocardial surface). The used sample points are depicted in Figure 6, bottom-left. For the computation of the arteriolar and capillary flows, we use the standard expressions:

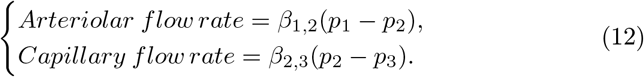

From the waveforms of Figure 6-top we can observe cyclic patterns of flow also in the microvasculature, similar to what obtained in the epicardial arteries (see Figure 5, left column) with wider oscillations going from the epicardium to the endocardium. These oscillations are lower in the capillary rather than arteriolar flow, suggesting a dampening effect of the microcirculation similar to what is observed in the peripheral circulation as a response to the pulsatility of the aortic pressure. This effect is observed to an increasing extent from the subendocardium to the subepicardium and it is also characterized by a time delay of the waveforms because of the compliance of the vessels wall.

**Figure 6:**
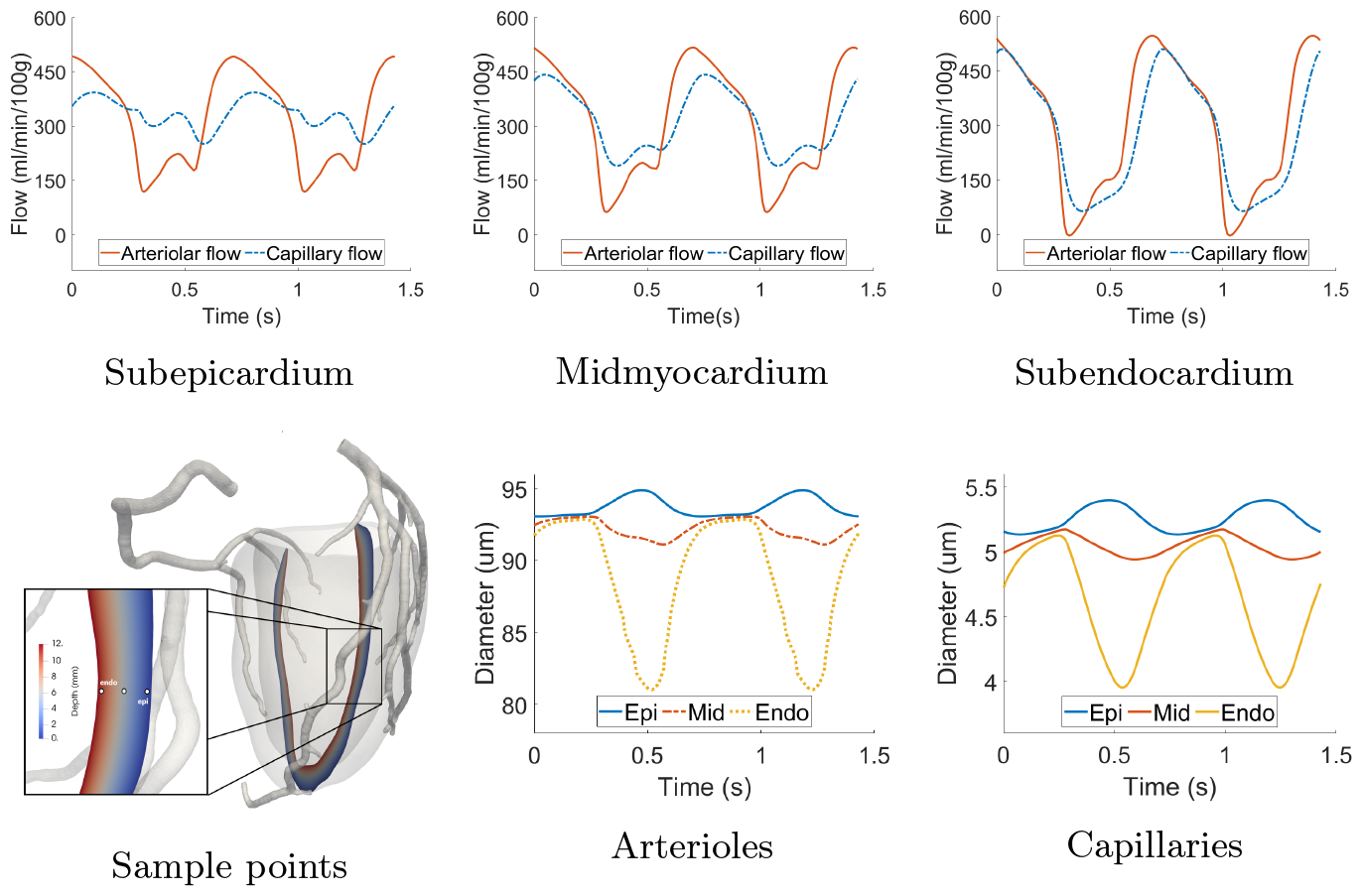
Top: arteriolar and capillary flow over time at three sample points in the mid-anterior wall at different depth locations. Bottom, middle and right: arteriolar and capillary diameters over time at the same sample points. Bottom, left: localization of the three sample points used for the flow and diameter computation. All results reported refer to the calibrated patient P1.

Figure 6, bottom-middle and bottom-right, reports the diameters of arterioles and capillaries computed from the pressures through relations (5)-(6) at the same sample points. Substantial differences are observed for both arterioles and capillaries at the various depths: slight increase in diameter at the subepicardium, slight decrease at the midmyocardium and marked decrease at the subendocarium. This behaviour, as well as the increasing flow oscillations, is consistent with the increase of the intramyocardial pressure from the epicardium to the endocardium and shows good agreement with experimental data of phasic diameter change in the arterioles (systolic/diastolic diameter change of ≃ −20% to ≃+2% from endocardium to epicardium, as reported in [30] from various animal measures).

Figure 7 reports the 3D results for coronary flow, coronary pressure and capillary flow at three key time instants over the cardiac cycle in the two patients. Consistently with the flow waveforms reported Figure 5 and 6, we observe that the diastolic flow is much higher than the systolic one both at the level of the capillaries and, in particular, of the epicardial arteries, despite the upstream pressure in the aorta being higher at the systolic peak. Also, we found a much higher pressure drop along the epicardial arteries at the early diastole (Δ*p* ≃9 − 15 mmHg) rather than at systolic peak (Δ*p* ≃ 4− 5 mmHg), which is a consequence of the higher diastolic flow. At the systolic onset (Figure 7-middle), the aortic pressure is at its lowest since the aortic valve is still closed; however, ventricular contraction is generating a high intramyocardial pressure which is the responsible of the inversion of the pressure gradient along the epicardial coronaries. This leads to a disturbed coronary flow (see representations in Figure 7) with vortexes at the proximal bifurcations, without never really establishing a full retrograde flow, most likely due to inertial effects and the brevity of this phase. Lastly, we observe that capillary flow distribution shows high regional heterogeneity in diastole, whereas it exhibits predominantly transmural heterogeneities in systole, when local hemodynamics is dominated by the intramyocardial pressure.

**Figure 7:**
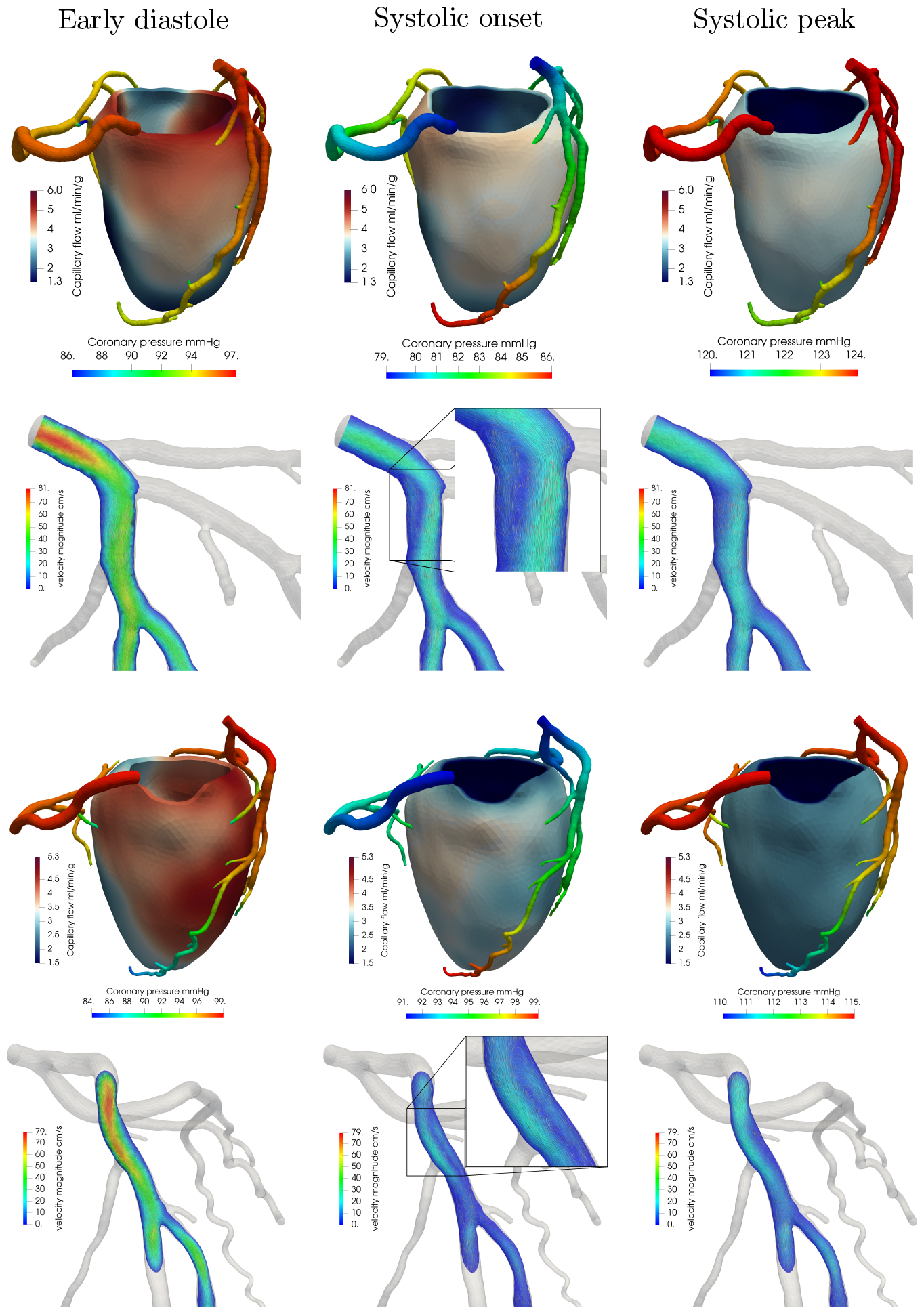
3D epicardial coronary pressure and capillary flow, computed as in the second line of 12 (top subpanel), and detail of the blood velocity in the left anterior descending artery (bottom subpanel), at three selected instants of the cardiac cycle. Notice that the scales for the coronary pressure are different at the three time instants to better highlight the key features. Patients P1 (top) and P2 (bottom).

Figure 8 reports the blood velocity at diastolic flow peak computed in the three main arteries, whereas Table 3 reports a summary of the most hemodynamically relevant quantities obtained from the simulations, for both patients P1-P2. The obtained velocities in the LAD and LCX (Left Circumflex) arteries are in good agreement with the *in vivo* Doppler measures reported in [44] for hyperemic conditions (48.8 ±14.3 *cm* · *s*^−1^ and 43.9 ±11.5 *cm* · *s*^−1^ for the LAD and LCX). Compared to these measures, our velocity results are on the lower side due to our two cases showing no anatomical lesions in the coronary arteries, while the data reported in [44] refer to a mixed population which includes also stenotic arteries where velocities may be much higher. In the case of the RCA, we observe an underestimation of blood velocity with respect to the measures (42.4 ± 12.4 *cm* · *s*^−1^) which is due to the absence, in our model, of the blood flow perfusing the right ventricle. We observe a significant inter-patient variability regarding the LAD-LCX flow subdivision, with values of 85%-15% for patient P1 and 62%-38% for patient P2, which may be related to the LAD/LCX diameter ratio (1.80 and 0.944 for patient P1 and P2, respectively) as well as to the myocardial mass perfused by each branch. Specifically, our patient P1 exhibits a high caliber first diagonal branch that originates from the proximal segment of the LAD and perfuses much of the territories normally perfused by the circumflex artery, which in this patient shows a much lower diameter. Conversely, in patient P2 the two branches have approximately the same caliber and a perfused myocardial mass much more balanced between them, resulting in a flow rate more evenly distributed. These results suggest that the specific anatomy plays a major role in flow subdivision and thus cannot be overlooked in computational frameworks that need an explicit prescription of such subdivision.

**Table 3:** Summary of hemodinamically relevant quantities computed for patients P1-P2. Velocities are averaged on the cross-section landmarks reported in Figure 8, diameters and flow rates are computed at the same landmarks.

| Artery | Quantity | Patient P1 | Patient P2 |
| --- | --- | --- | --- |
| LAD | Cross-section avg. peak velocity $cm \cdot s^{-1}$ | 39 | 44 |
| | Flow rate $ml \cdot min^{-1}$ | 258 | 110 |
| | Diameter $mm$ | 5.4 | 3.4 |
| LCX | Cross-section avg. peak velocity $cm \cdot s^{-1}$ | 32 | 36 |
| | Flow rate $ml \cdot min^{-1}$ | 44 | 66 |
| | Diameter $mm$ | 3.0 | 3.6 |
| RCA | Cross-section avg. peak velocity $cm \cdot s^{-1}$ | 19 | 16 |
| | Flow rate $ml \cdot min^{-1}$ | 105 | 60 |
| | Diameter $mm$ | 4.8 | 4.1 |

**Figure 8:**
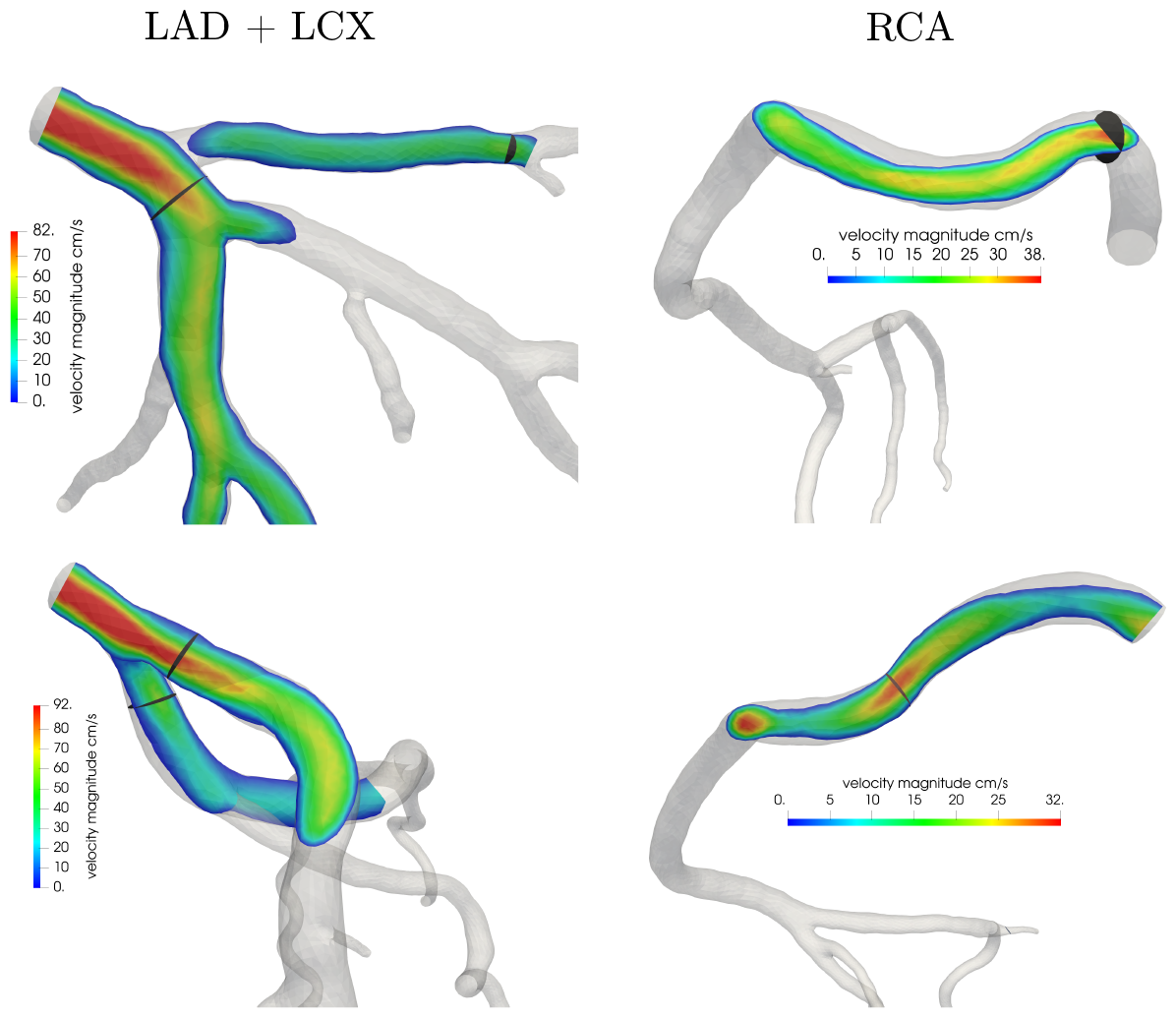
Velocity magnitude at the diastolic flow peak for the main artery branches: LAD, LCX artery, RCA for patient P1 (top) and P2 (bottom), with indication of the cross-section used to find the peak velocity values. All the cross sections are placed in the most proximal segments of the corresponding artery, i.e. segment 1 for RCA, segment 6 for LAD and segment 11 for LCX.

### 3.2 Parameter sensitivity analysis

All the results presented in Section 3.1 are obtained using literature values for vessels length densities (*nL*)_*i*_ and values of specific permeability *κ* computed through eq. (7) so that the resulting Darcy permeabilities *K*_*i*_ were in line with previous computational studies [8, 9]. Since inter-patient variability of these parameters could be relevant for predictive applications, we report here a sensitivity analysis performed to quantify how and to what extent these parameters affect the results.

Figure 9 reports the results of the sensitivity analysis on the specific permeability *κ* in the range (5 ·10^−10^ −5· 10^−8^) *m*^−1^. This affects the Darcy permeabilities *K*_*i*_ (see (7)). For all the three values considered, we observe no relevant changes in the waveforms of arterial flow and in-space average arteriolar pressure over time (top row) and in the overall time-averaged capillary flow (MBF, bottom row). Regarding the latter quantity, we notice an increased heterogeneity in MBF regional distribution as *κ* decreases. This is a consequence of the piecewise constant mass source term, among the perfusion regions, in the first compartment of the Darcy model (see (4)), representing blood flow in the different perfusion regions coming from the associated feeding arteries. However, the pressure and the resulting MBF is smoothed by the Darcy permeabilities *K*_*i*_, with increasing smoothing effect among regions for higher values of *κ*. Physically, this means that *κ* regulates how independent each perfusion territory is from the surrounding ones, with lower values associated with a more compartmentalized perfusion, without significantly affecting the average value. Since the range of values analyzed is quite large (two orders of magnitude), we conclude that our model, in terms of in-space average quantities, has a rather low sensitivity towards *κ*.

**Figure 9:**
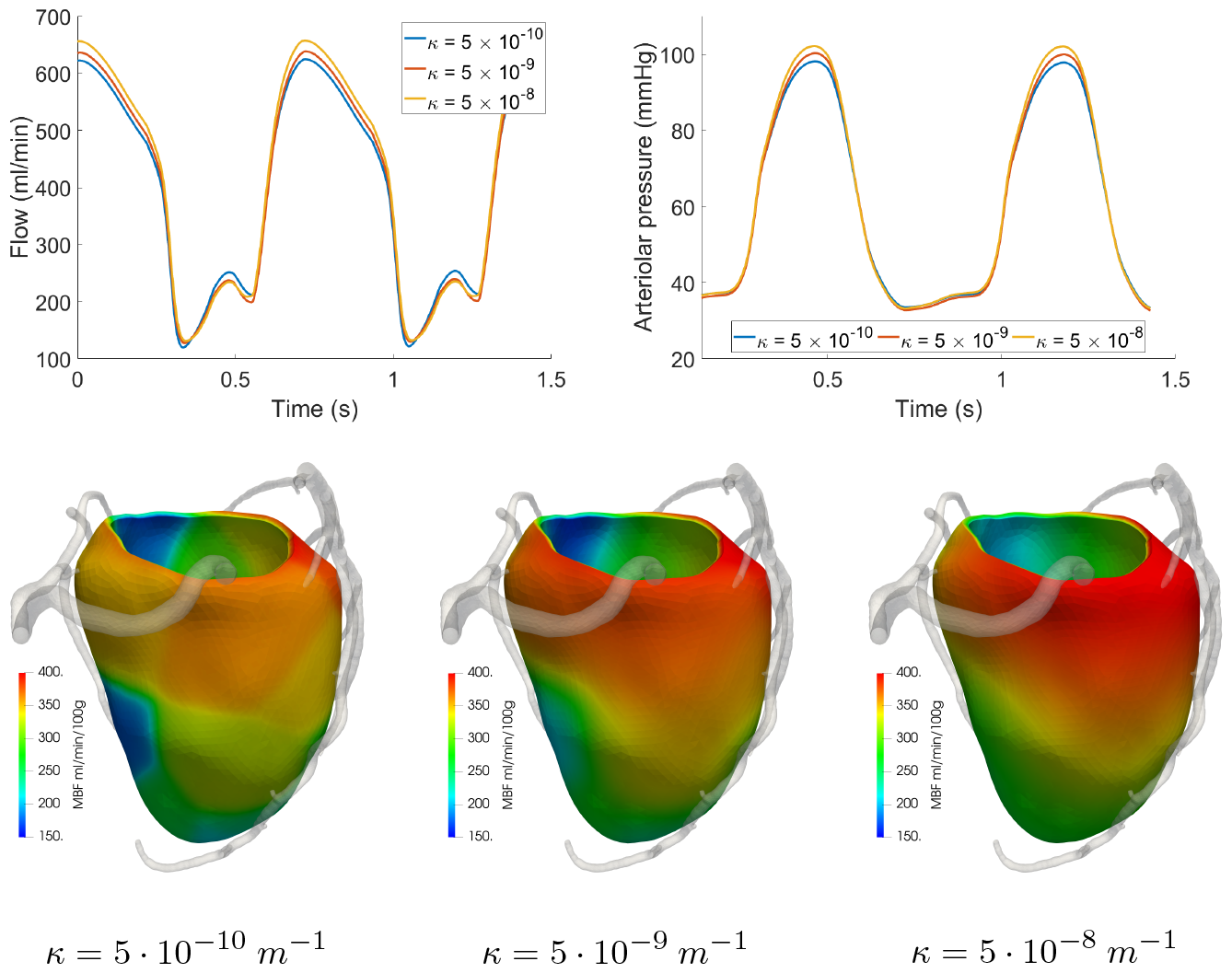
Effect of *κ* in the range (5 ·10^−10^ −5 ·10^−8^) *m*^−1^. Top: total arterial inflow (left) and average in-space arteriolar pressure (right) waveform over time. Bottom: 3D distribution of time-averaged capillary flow (MBF) over the myocardium.

Figure 10 reports the results of the sensitivity analysis on the vessels length density (*nL*)_*i*_ when the values associated to each Darcy compartment are scaled all by the same factor, in the range 0.75x 2x w.r.t. the values reported in Table 1. From the waveforms in Figure 10-top, we can see that the coronary flow and arteriolar pressure show a high sensitivity towards these parameters, which is not surprising given the direct proportionality between them and the inter-compartment Darcy conductances *β*_*i,j*_ (see (9)). We notice also that the systolic flow is much less affected than the diastolic one, which is likely due to the fact that vessels length densities (*nL*)_*i*_ affect also the compliance term in (4), leading to the balance of opposite contributions. Since diastolic flow is instead highly dependent on (*nL*)_*i*_, we observe an overall logarithmic dependence of in-space average MBF vs (*nL*)_*i*_, with the analytical fitted relationship reported in Figure 10, bottom-right. Other effects include higher arteriolar pressures (higher conductances in the Darcy model means that there is a higher pressure jump between the capillary compartment and the veins) and negligible effects on MBF distribution, which is mostly regulated by Darcy permeabilities *K*_*i*_. Even if *K*_*i*_ are directly proportional to (*nL*)_*i*_ (see (7)), the range analyzed for this parameter is too narrow, compared to the sensitivity we found for *κ*, to lead to relevant differences.

**Figure 10:**
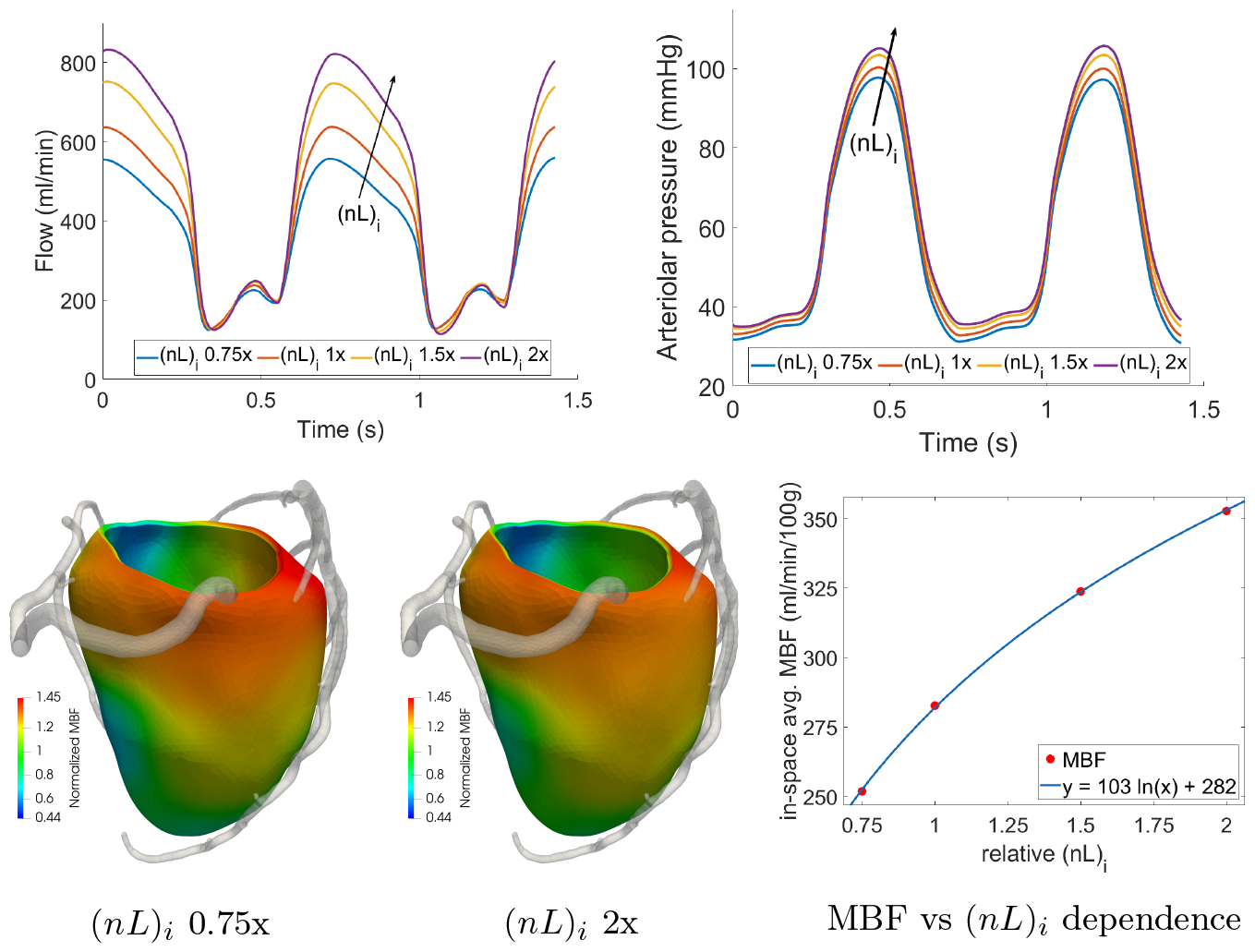
Effect of (*nL*)_*i*_ values scaled homogeneously in the range 0.75x 2x w.r.t. the values reported in Table 1. Top: waveforms over time of total arterial inflow (left) and arteriolar pressure (right). Bottom left: 3D distribution of time-averaged capillary flow (MBF) normalized over the mean value to highlight the distribution; bottom right: logarithmic fit built on the dependence of in-space averaged MBF vs (*nL*)_*i*_.

Since the model may show different responses to variations in single compartment length densities, for example due to the different constitutive relationships implemented for arterioles and capillaries, we also run simulations where (*nL*)_*i*_ is changed non-uniformly in the three compartments. Figure 11 reports the results in terms of total arterial inflow and pressure in small arterioles and arterioles in the two following scenarios (base refers to Tab. 1),:

1. (*nL*)_1_ doubled (w.r.t. base), (*nL*)_2_ halved, (*nL*)_3_ base,
2. (*nL*)_1_ doubled, (*nL*)_2_ base, (*nL*)_3_ halved;

We can see that the flow waveform is predominantly affected by the capillary length density, with an increased oscillation in systolic flow and a slight decrease in diastolic flow in scenario 2. Pressure in the small arteries relatively unaffected in both scenarios, whereas diastolic arteriolar pressure shows a moderate (≃5 mmHg) and relevant (≃10 mmHg) increase in scenarios 1 and 2, respectively.

**Figure 11:**
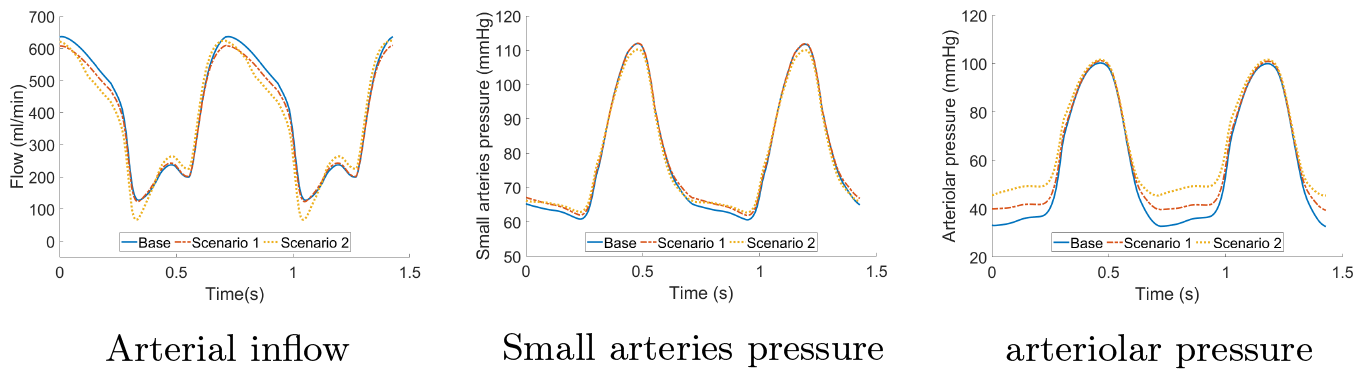
Effect of different combinations of (*nL*)_*i*_ values on flow and pressure waveforms. The Base scenario refers to the parameters reported in Table 1, whereas the other two scenarios represent relative scaling (as indicated) with respect to the base parameters.

From this analysis on the changes of the most relevant parameters, we can conclude that:

1. *κ* mainly regulates the sharpness in the MBF gradients between adjacent perfusion regions. Higher values of *κ* lead to smoother gradients and a more homogenized perfusion.
2. Parameters (*nL*)_*i*_ regulate the pressure jumps between compartment *i* and its adjacents. We also observe that (*nL*)_*i*_ has a relevant effect on diastolic flow, which greatly increases as (*nL*)_*i*_ increase, while its effects on systolic flow is negligible. Out of all the Darcy compartments, the capillary length density (*nL*)_3_ shows the highest impact on the results.

## 4 Discussion

Precise and effective modeling of coronary hemodynamics and cardiac perfusion is a daunting task due to the complexity of physical phenomena occurring during the heartbeat. The multiscale nature of the coronary circulation and the presence of a cyclic mechanical activation of the muscle represent key features that have a deep impact on the hemodynamics at the different scales. Several clinical studies demonstrated that the quantification of blood flow at the capillary level and over the various regions of the cardiac muscle adds significant prognostic value in the management of patients suffering from coronary artery disease [1, 2, 45, 46]. In these studies, *in vivo* functional imaging techniques such as nuclear imaging (Positron Emission Tomography - PET, Single Photon Emission Computed Tomography - SPECT) or stress CTP are used to extract a 3D map of myocardial perfusion based on the radiotracer signal (in the nuclear imaging tests) or the time-attenuation curves of the contrast agent over the heartbeat. In all the cases, the myocardial blood flow is quantified using both diastolic and systolic flow values, so the development of models able to accurately capture both the diastolic and systolic 3D microcirculatory hemodynamics is of paramount importance.

In this work, we present a fully distributed, 3D mathematical model of coronary hemodynamics from the epicardial arteries (Navier-Stokes formulation) to the microcirculation (multi-compartment Darcy formulation), that also includes for the first time the compliance of the microvessels and the presence of a cyclic external pressure representing the effects of cardiac contraction. In the model, we include nonlinear constitutive relationships for microvessels compliance that we built on experimental data and we propose a new formulation for the Darcy parameters based on histologically relevant quantities, e.g. the local length density of vessels. This aspect is particularly relevant since such direct link can be exploited for a robust and precise calibration of the model, potentially tuning these parameters among the various myocardial regions, for example to distinguish between more and less vascularized territories.

Our model, applied to the simulation of hyperemic coronary flow in real clinical cases with geometries segmented from CT images, is successful in reproducing the phasic coronary flow pattern of high diastolic arterial inflow and systolic venous outflow, showing excellent agreement with experimental literature data with respect to the shape of the flow and pressure curves, the time evolution of diameters of microvessels and the differences at the various depths in the cardiac muscle. Notice that the calibration of the Darcy parameters seems to be robust with respect to the patients. Indeed, we used only limited information about P1 hemodynamics (namely the in-space average MBF) for the calibration and this led to consistent results also for patient P2.

From a clinical perspective, our study provides an important contribution in view of the development of a tool to predict the whole hemodynamics in the coronary tree. To this regard, in [21] we have shown that our model is successful in predicting Fractional Flow Reserve (FFR), along the lines of previous works [47–51], and in providing information on global Myocardial Blood Flow (MBF). Here, we instead focus on microcirculation, providing information on hemodynamics also at the tissue level. With this aim in mind, we remark that our model specifically adopts a fully distributed description of microvasculature hemodynamics and over the whole heartbeat, making it possible to quantify regional and transmural distribution of MBF over the myocardium. This will allow us in principle to include in our model the specific effects of other pathologies such as left ventricular hypertrophy and scenarios of altered contractility.

In the direction of developing a tool which is able to predict also MBF distribution, we present here some further (preliminary) results on the comparison of our simulated MBF maps with the outcome of the stress-CTP imaging exam performed on the same two patients: Figure 12 reports such comparison, where the compared quantity is the MBF averaged over each perfusion region for patients P1 and P2.

**Figure 12:**
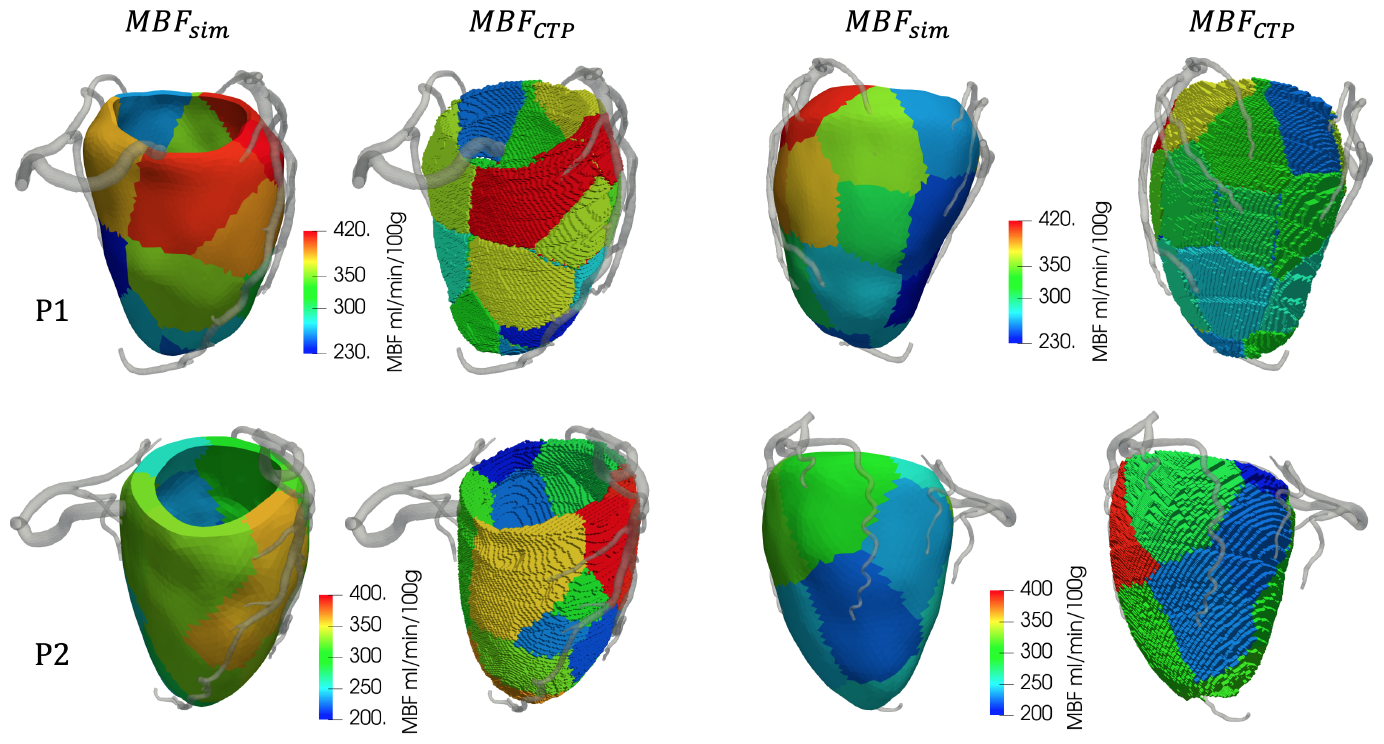
Comparison between region-averaged MBF from our simulations (*MBF*_*sim*_) vs region-averaged MBF from stress-CTP imaging (*MBF*_*CTP*_). Top: patient P1; Bottom: patient P2. Left block: anterior view; Right block: posterolateral view.

We can see from these results that our model is able to reproduce the MBF distribution in both the analyzed patients, obtaining values in good accordance with the clinical perfusion maps. Notably, we did not observe any region with *MBF <* 150 ml/min/100g, which is the critical threshold for the detection of regions susceptible to inducible ischemia. This was in accordance with the outcomes of the stress-CTP exam, thus we do not report any false-positive region in the two patients, despite the high heterogeneity in MBF values across the various regions.

This work presents some limitations. Firstly, we are aware that two patients are not enough to conclude that our model can consistently predict the absence of perfusion defects in a given patient, i.e. assess the model specificity in a robust way. However, this is not a statistical study, rather a mechanistic one aimed at predicting hemodynamics, so that the number of patients is necessarily not high. Also, even though the analyzed subjects were symptomatic patients (as specified in Section 2.7), we did not examine any subjects with diagnosis coronary artery disease, so we cannot assess the sensitivity of the model in the prediction of the presence and localization of perfusion defects. Further studies applied to a large and mixed population would be highly desirable to these aims. Still, we believe that the good agreement we found in the MBF distribution between our simulations and the clinical maps (see Figure 12) represents a very strong starting point in this direction.

Other limitations regard modeling choices and clinical applicability:

1. We considered constant values along the whole vasculature (i.e. independent of the Darcy compartment) for both the viscosity *μ* and the specific permeability *κ*, see (7). These choices have been motivated by noticing that both fluid-matrix interaction and rheology, although they may change along the vasculature, do not have an influence,in our compliant model, in recovering the right qualitative behaviours of the quantities of interest (phasic flow, systolic impediment, etc.). However, we believe that a deeper investigation of the quantitative influence of these parameters should be in order for future studies;
2. We did not consider regional nor transmural differences in the vessels properties. For example, it has been found that capillary density and diameter are higher at the subendocardium rather than at the subepicardium [52] but it is not clear how this could impact the MBF results in physiological and pathological scenarios;
3. Our model does not take into account that the intramural vessels (small arteries and microvessels) have a precise course into the cardiac muscle, that is a transmural course with increasing ramifications moving from the epicardium to the endocardium. The main implication of this, which is not captured by our model, is that subendocardial vessels are in general in a downstream position with respect to subepicardial vessels of the same Darcy compartment, thus their intraluminal pressure is likely different, with lower values in the diastolic phase;
4. Regarding clinical applicability, an extended analysis with validation on a large and mixed population should be in order. This would require to strongly address the issue of the model calibration, potentially including space-dependent and personalized Darcy parameters. For a predictive application where the use of data coming from stress-CTP imaging is not an option, this calibration should use only geometrical information, basic patients metadata or data coming from first-pass exams such as rest coronary CT angiography.

## Acronyms

*CAD*: Coronary Artery Disease
*CBF*: Coronary Blood Flow
*cCTA*: coronary Computed Tomographic Angiography
*FFR*: Fractional Flow Reserve *LAD* Left Anterior Descending
*MBF*: Myocardial Blood Flow
NS: Navier-Stokes
*RCA*: Right Coronary Artery
*stress-CTP*: stress Computed Tomographic Perfusion

## List of Symbols

*p*_*i*_: Pore pressure in Darcy compartment *i K*_*i*_ Permeability of Darcy compartment *i*
*φ*_*i*_: Fluid volume fraction of Darcy compartment *i*
*β*_*i,j*_: Inter-compartment conductance between Darcy compartments *i* and *j P*_*LV*_ Pressure in the left ventricular chamber
*P*_*im*_: Intramyocardial pressure
*θ*_*i*_: Mass source/sink term in Darcy compartment *i γ* Conductance of venous circulation
*p*_*ra*_: Right atrium pressure
(*nL*)_*i*_: Vessels length density of Darcy compartment *i*
*A*_*i*_: Average cross section of vessels in Darcy compartment *i*
*C*_*i*_: Compliance of vessels in Darcy compartment *i*
*μ*: Blood dynamic viscosity
*κ*: Specific permeability of cardiac tissue to blood flow
*δ*_*i*_: Morphometry conductance factor of Darcy compartment *i*

## Acknowledgements

GMP, FR, SB, CV are members of the INdAM group GNCS “Gruppo Nazionale per il Calcolo Scientifico” (National Group for Scientific Computing). CV has been partially supported by the Italian Ministry of University and Research (MIUR) within the PRIN (Research projects of relevant national interest) MIUR PRIN22-PNRR n. P20223KSS2 “Machine learning for fluid-structure interaction in cardiovascular problems: efficient solutions, model reduction, inverse problems, and by the Italian Ministry of Health within the PNC PROGETTO HUB LIFE SCIENCE - DIAGNOS- TICA AVANZATA (HLS-DA) “INNOVA”, PNC-E3-2022-23683266–CUP: D43C22004930001, within the “Piano Nazionale Complementare Ecosistema Innovativo della Salute” - Codice univoco investimento: PNC-E3-2022- 23683266.

## Funding

This work has been supported by Bracco Imaging S.p.A., by Consiglio Nazionale delle Ricerche (CNR) and by Italian PNRR research funding, Missione 4, DM226/2021.

## Notes

### Competing Interest Statement

The authors have declared no competing interest.

